# Genetic and gene expression analysis of flowering time regulation by light quality in lentil

**DOI:** 10.1101/2021.02.14.429948

**Authors:** Hai Ying Yuan, Carolyn T. Caron, Larissa Ramsay, Richard Fratini, Marcelino Pérez de la Vega, Albert Vandenberg, James L. Weller, Kirstin E. Bett

## Abstract

Flowering time is important due to its roles in adaptation to different environments and subsequent formation of crop yield. Changes in light quality affect a range of developmental processes including flowering time, however little is known about light quality induced flowering time control in lentil. This study aims to investigate the genetic basis for differences in flowering response to light quality in lentil.

We explored variation in flowering time caused by changes in red/far-red related light quality environments of a lentil interspecific recombinant inbred line population developed from a cross between *Lens culinaris* cv. Lupa and *L. orientalis* accession BGE 016880. A genetic linkage map was constructed and then used for identifying QTL associated with flowering time regulation under different light quality environments. Differential gene expression analysis through transcriptomic study and RT-qPCR were used to identify potential candidate genes.

QTL mapping located 13 QTLs controlling flower time under different light quality environments, with phenotypic variance explained ranging from 1.7 to 62.9%. Transcriptomic profiling and gene expression analysis for both parents of this interspecific RIL population identified flowering-related genes showing environment-specific differential expression (flowering DEGs). One of these, a member of the florigen gene family FTa1 (*LcFTa1*) was located close to 3 major QTLs. Furthermore, gene expression results suggests two other florigen genes (*LcFTb1* and *LcFTb2*), MADS-box transcription factors like *LcAGL6/13d, LcSVPb, LcSOC1b* and *LcFULb*, as well as bHLH transcription factor *LcPIF6* and Gibberellin 20 oxidase *LcGA20oxC,G*, may be involved in the light quality response as well.

Our results show that a major component of flowering time sensitivity to light quality is tightly linked to *LcFTa1* and associated with changes in its expression. This work provides a foundation for crop improvement of lentil with better adaptation to variable light environments.

## INTRODUCTION

Flowering, a transition from vegetative growth to reproductive growth is one of the most important events for flowering plants. The switch from vegetative to reproductive growth is essential for crop production when seeds or fruits are to be the end products. Plants need to recognize and process a wide range of environmental and internal cues, and then consolidate these into a single effective developmental choice: to flower or not (Putterill *et al*., 2004). Some of the main environmental factors that control flowering time include photoperiod and temperature, but light quality is also known to have an important influence (Adams *et al*., 2009; Casal, 2013), and acts as a crucial signal that communicates the close presence of neighbouring plants (Adams *et al*., 2009). One of the main components of this signal is a reduction in the red to far-red ratio (R/FR) which results from the selective absorption of red wavelengths by chlorophyll (Smith and Whitelam, 1997). This reduction in R/FR is primarily sensed through the phytochrome family of photoreceptors, which mediate its effects on a wide range of developmental processes, including increased stem elongation, reduced branching and accelerated flowering (Smith and Whitelam, 1997; Quail, 2002; Kami *et al*., 2010).

Studies of flowering time control have identified numerous flowering time genes and defined genetic regulatory networks in model species (Song *et al*., 2015; Hori *et al*., 2016; Gol *et al*., 2017), as well as in some legumes (Weller and Ortega, 2015; Ridge *et al*., 2017; Ortega *et al*., 2019; Lin *et al*., 2020). Many of these genes contribute to the natural genetic variation for flowering time and for related growth and yield traits, and provide adaptation of crop plants to various locations and management practices (Jung and Müller, 2009; Bouchet *et al*., 2013; BlüMel *et al*., 2015). The importance and genetic basis of variation in response to light quality specifically is relatively poorly understood in many crop species, although many of the genes defined in Arabidopsis are widely conserved (Hecht *et al*., 2005; Adams *et al*., 2009; Leijten *et al*., 2018).

A genetic understanding of flowering time regulation is an important general objective in plant breeding, as it facilitates the generation of new varieties that will be better adapted to a specific environment. Genetic analysis of flowering time has been conducted in many legume crops (Pérez-Vega *et al*., 2010; Cruz-Izquierdo *et al*., 2012; Upadhyaya *et al*., 2015; Liu *et al*., 2016; Ridge *et al*., 2017), including lentil (Sarker *et al*., 1999; Fratini *et al*., 2007; Tullu *et al*., 2008; Fedoruk *et al*., 2013). However, few studies so far have investigated the genetic basis for flowering responses to light quality, and none in lentil.

Cultivated lentil (*Lens culinaris* Medik.) is the third most important cool-season grain legume (FAO, 2015). Lentils have high nutritional and health benefits, and like other crop legumes, play a significant role in supporting environmentally sustainable agriculture due to their nitrogen fixation ability. In addition to the cultivated lentil, there are six wild species within the genus Lens (Van Oss *et al*., 1997) and they contain rich genetic diversity that can be used for genetic improvement of cultivated lentils (Vail *et al*., 2011; Podder *et al*., 2013; Bhadauria *et al*., 2017). In a recent study, we found that the flowering time of most wild lentil genotypes was not significantly affected by light quality changes, whereas it was consistently accelerated under the low R/FR conditions in cultivated lentil (Yuan *et al*., 2017). This variation in flowering time sensitivity toward the light quality change indicated that genes or specific alleles associated with this trait could be used to select or modify flowering time in cultivated lentil.

In this study, we used an interspecific recombinant inbred line (RIL) population of lentil (Fratini *et al*., 2007) whose parents had contrasting flowering responses to light quality change. We characterized the variations in flowering responses of individual RILs to contrasting R/FR environments and used a high-density genetic linkage map to identify QTLs associated with differential sensitivity to light quality. *De novo* transcriptomic analysis was used to characterize the effects of light quality on gene expression, and together with the newly available *L. culinaris* genome (CDC Redberry, v 2.0) (Ramsay *et al*., 2019), hereby referred to as the reference genome, we were able to identify and evaluate potential candidate genes for this important response.

## MATERIALS AND METHODS

### Plant material, growth conditions and phenotypic evaluation

An interspecific recombinant inbred line (RIL) population developed from a cross between *L. culinaris* cv. Lupa and *L. orientalis* accession BGE 016880 (Fratini *et al*., 2007) was used in this study. Both parents have contrasting flowering time responses to light quality change, with the wild parent being less sensitive (Yuan *et al*., 2017). The RIL population of 93 individuals and parents were grown and evaluated in two Conviron GR178 walk-in plant growth chambers with contrasting R/FR ratios, but similar light quantities based on the photosynthetically active radiation (PAR). The high R/FR light quality condition was the natural condition in the growth chamber fitted with T5 835 High Output Fluorescence bulbs (Philips, Andover, MA, USA). The low R/FR light quality condition was reached by adding evenly spaced PfrSpec™ LED light panels (Fluence Bioengineering, Inc., model RAY44, peak spectrum at 730nm, Austin, Texas, USA) into the light bank fitted with the T5 835 High Output fluorescent bulbs. The spectral photon flux and PAR of each light condition was measured using a spectroradiometer (Apogee Instruments, Model PS-300, Logan, UT, USA). The R and FR values were calculated using the spectral photon flux at 650–670 and 720–740 nm, respectively (Smith, 1982). The high R/FR condition had an R/FR ratio of 7.30 ± 0.14 with PAR at 402.2 ± 33.6 µmol/m^2^s, and the low R/FR condition had an R/FR ratio of 0.19 ± 0.01 with a PAR level very close to the high R/FR condition at 395.6 ± 32.9 µmol/m^2^s. The spectral distribution of these two light conditions is shown in **Supplementary file S1_Fig. S1A**.

Prior to planting, seeds of all RILs and parents were stored at −20 °C for one week and then the seed coats were nicked to improve imbibition and germination. Square, 10 cm pots were filled with growth medium consisting of 50% Sunshine Mix #3 and 50% Sunshine Mix #4 (Sun Gro Horticulture Canada Ltd., Seba Beach, AB Canada). Two seeds were sown in each pot and thinned to a single plant after emergence. The RILs and the parents were planted in a completely randomized design with four technical replicates per genotype under each light quality condition and the experiment was repeated once. Plants were grown and maintained at 22 °C/16 h day and 16 °C/8 h night for both conditions and repeats. Days to flower (DTF) were calculated based on days from emergence to R1 (one open flower at any node) (Erskine *et al*., 1990). Levene’s test of homogeneity of variance was done for days to flower using R (R Core Team, 2013). There were no significant effects of repeat, so data were averaged over the two repeats. For each individual the trait “Flowering Time Sensitivity” (FTS) was calculated as the ratio of the difference in DTF between two conditions divided by the sum of the DTF under two conditions so as to avoid bias due to the underlying differences in DTF.

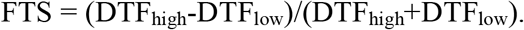

### RADseq library preparation, sequencing and SNP calling

Illumina sequencing libraries were prepared using genomic DNA from the RIL population for restriction site-associated DNA sequencing (RADseq). HindIII and NlaIII were chosen as two restriction enzymes to digest the genomic DNA. Detailed library preparation procedures were described in von Wettberg et al. (von Wettberg *et al*., 2018). Fragments were sequenced as single-end 100-bp reads on an Illumina HiSeq4000 at the University of California at Davis Genome Core Facility. Reads were mapped to the *L. culinaris* v2.0 reference genome using Bowtie allowing only end-to-end matches (Langmead and Salzberg, 2012). Variant calling was performed with Samtools (Li *et al*., 2009) and output in VCF format (Danecek *et al*., 2011). SNP Variants were filtered using VCFtools (Danecek *et al*., 2011) to exclude sites that contain an indel, keep variants that have no more than 66% missing data with a minimum quality score of 30, and include only bi-allelic sites. After filtering, SNP variants were converted to ABH format, markers that were 100% identical in sequential order along the chromosome (including missing) were binned and further filtered for markers where only the parents were different. Initially, 84,721 SNPs were generated from the RIL population and after filtering and binning, 15,686 SNPs were retained for linkage mapping.

### Linkage map construction and QTL mapping

Linkage map construction was performed with IciMapping software (Meng *et al*., 2015). BIN functionality within the software was used to remove redundant SNP markers that have identical segregation in the RIL population. After binning, SNP markers were grouped using logarithm of the odds (LOD) threshold value of 8.0. Linkage groups (LGs) were assigned using the genomic position of SNP markers on the *L. culinaris* v2.0 reference genome (Ramsay *et al*., 2019). The REcombination Counting and ORDering (RECORD) algorithm (Van Os *et al*., 2005) was used to order the SNPs within each LG. The Kosambi mapping function (Kosambi, 1943) was used to convert the recombination fractions into additive genetic distance (centiMorgans). Rippling of the SNP markers was performed by permutation of a window of 8 markers and using sum of adjacent recombination frequencies (SARF) as rippling criterion. The linkage map was further adjusted using the R/qtl package (Broman *et al*., 2003). The marker order within a linkage group that gave the shortest genetic distance was chosen as the final map.

QTL mapping was performed using BIP functionality in IciMapping software (Meng *et al*., 2015). For QTL identification, the genotyping data was integrated with the phenotypic data. Inclusive Composite Interval Mapping of Additive (ICIM-ADD) was used to detect additive QTL. The walking speed chosen for QTL mapping was 1 cM. A 1000 permutation test was applied to decide the logarithm of odds (LOD) thresholds (P ≤ 0.05) to determine the significance of identified QTLs. A One-LOD support interval was calculated for each QTL to obtain a 95 % confidence interval. The percentage of phenotypic variance explained by each QTL in proportion to the total phenotypic variance was estimated and QTLs with a positive or negative additive effect for the trait imply that the increase in the phenotypic value of the trait is contributed by alleles from *L. culinaris* cv. Lupa or *L. orientalis* BGE 016880, respectively. QTL for all traits were named according to McCouch et al. (1997) and alphabetical order was used for QTLs on the same linkage group. Linkage map was drawn using LinkageMapView package in R (Ouellette *et al*., 2018).

### Transcriptomic analysis and RT-qPCR verification of the candidate genes

Both *L. culinaris* cv. Lupa and *L. orientalis* BGE 016880 were grown under two different R/FR ratio environments, same as the ones used for RIL population screening. Leaf sample collections started two weeks after emergence (T1 stage) and continued once a week for five weeks (T2 to T5 stage). T1 sampling stage was two weeks before *L. orientalis* BGE 016880 flowered under low R/FR environment and **Supplementary file S1_Figure S2** shows relations between the sampling stages and the flowering times for both BGE 016880 and Lupa under different R/FR environments. Samples were taken at the same time of the day for each collection. Each biological replicate consisted of leaf material collected from three individual plants and three biological replicates were used.

Total RNA was isolated using RNeasy Plant Mini Kit (QIAGEN, Germantown, MD, USA) according to the protocol provided with the kit. On-column DNase digestion was performed during the isolation process according to the kit instructions. Extracted RNA was quantified and qualified using a NanoDrop 8000 UV–Vis spectrophotometer (NanoDrop, Wilmington, DE, USA) and an Agilent 2100 Bioanalyzer (Agilent Technologies, Santa Clara, CA, USA) using an Agilent RNA 6000 Nano Assay. RNAseq libraries were constructed using Illumina TruSeq Stranded mRNA Sample Preparation Kit (Illumina Inc., San Diego, CA, USA) according to the protocol, and pooled libraries of 20 barcoded samples were sent for pair-end sequencing using an Illumina HiSeqTM 2500 sequencing platform (Illumina Inc., San Diego, CA, USA).

Qualities of the raw reads were checked using FASTQC (Andrews, 2010) and adaptor sequences were removed using TRIMMOMATIC (Bolger *et al*., 2014). Trimmed-reads were used for *de novo* assembly of transcriptome using Trinity (Grabherr *et al*., 2011). Salmon (Patro *et al*., 2017) was used to quantify the abundance of transcripts and 3D RNA-seq pipeline (Calixto *et al*., 2018; Guo *et al*., 2019) was used for the analysis of differential gene expression within species. A detailed method regarding differential gene expression analysis is described in **Supplementary file S1_Supplmentary Methods**. Genes with FDR adjusted p-values < 0.05 and |log_2_FC|≥1 were considered differentially expressed genes (DEGs; FC, Fold change).

A list of 244 lentil genes that showed high homology with genes involved in flowering in *Arabidopsis thaliana* (https://www.mpipz.mpg.de/14637/Arabidopsis_flowering_genes and (Higgins *et al*., 2010) and *Medicago truncatula* (Hecht *et al*., 2005; Putterill *et al*., 2013) was curated (**Supplementary File S2**) using in-house BLASTn search (https://knowpulse.usask.ca/blast/nucleotide/nucleotide) against the *L. culinaris* cv. CDC Redberry v2.0 reference genome. An e-value cut-off of 1e-5 or matched gene annotation in the lentil genome assembly was used to select the putative homologue. This list was used to explore the expression of known flowering genes in the lentil genome.

To confirm the reliability of RNAseq results, representative flowering-related genes were chosen for RT-qPCR using FAST SYBR® Green qPCR Master Mix (Applied Biosystems, Foster City, CA, USA) with a BioRad Real-Time PCR system. Actin (Lcu.2RBY.L011470) was selected as the reference gene. To derive the relative expression value, the delta-delta CT method was adopted (Livak and Schmittgen, 2001) and samples from T1 stage at low R/FR light quality environment were used as reference samples. The sequences of the primers used in qPCR were listed in supplemental information (**Supplementary file S1_Table S1**).

### Data availability

The RAD-seq data of the RIL population plus two parents (*L. culinaris* cv. Lupa and *L. orientalis* BGE 016880) as well as the raw RNAseq data from *L. culinaris* cv. Lupa and *L. orientalis* BGE 016880 have been submitted to the NCBI Sequence Read Archive with the IDs as PRJNA693555 and PRJNA693582. Related tables, figures and methods are given in **Supplementary file** S1. All other related data as well as the in-house developed scripts are given in **Supplementary files S2 to S6**.

## RESULTS

### Variation in flowering time response to light quality

To explore the genetic control of differences in flowering response to light quality we previously described for wild and domesticated lentil accessions (Yuan *et al*., 2017), we used a RIL population developed from an interspecific cross between *L. culinaris* cv. Lupa and an accession (BGE 016880) of *L. orientalis*, the putative progenitor of cultivated lentil (Mayer and Soltis, 1994). Flowering time in this population was assessed under two light environments with equivalent PAR but differing in R/FR (**Supplementary file S1_Fig. S1A)**. We also expressed the response of each line to light quality as a “flowering time sensitivity” index (FTS), which was calculated as (DTF_high_-DTF_low_)/(DTF_high_+DTF_low_), in order to avoid bias due to the underlying differences in DTF.

The wild parent, BGE 016880, flowered significantly earlier than the cultivated parent, Lupa, under both light quality environments and the wild parent had a low FTS compared to the cultivated parent (P < 0.05; **Figure 1** and **Supplementary file S1_ Fig. S1**). The RIL population segregated for days to flowering and for sensitivity to change in light quality, with probability density of the distributions centered on the mid-parent value under both light conditions and transgressing the parental range in both directions (**Fig. 1)**. Some indication of a bimodal distribution was evident for flowering under high R/FR and for flowering time sensitivity (**Fig. 1**), suggesting the possibility of a substantial contribution from a single major locus.

**Fig. 1.**
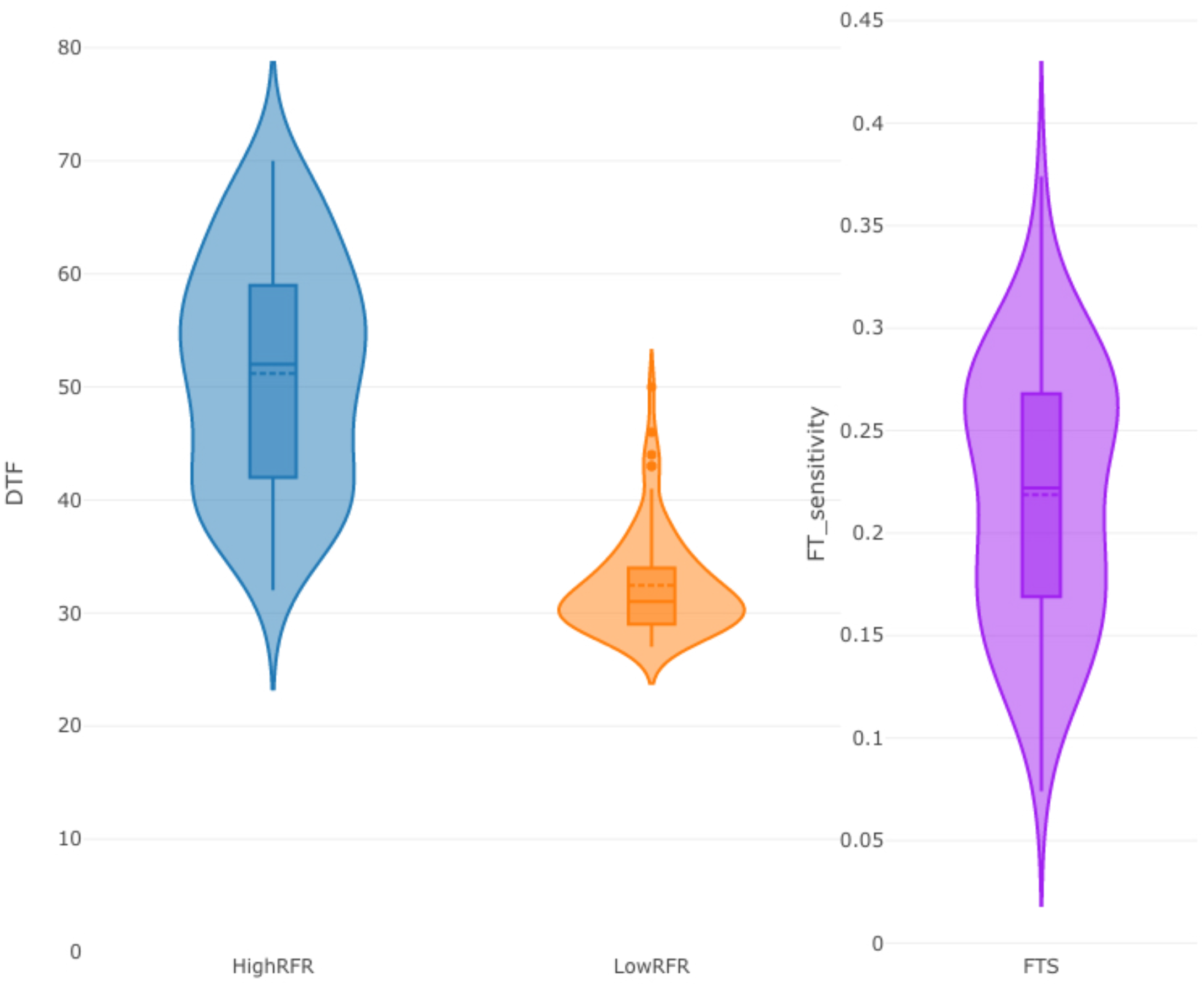
Days to flower of individuals from a Lentil interspecific RIL population developed from a cross between *L. culinaris* cv. Lupa and *L. orientalis* BGE 016880, grown under light quality environments differing in red to far-red ratio (R/FR) (left & middle) and flowering time sensitivity to changes in R/FR light quality environment (right). The violin plot outline illustrates kernel probability density and the width of the shaded area represents the proportion of the data located there. The inner section of the violin plot shows the box plot indicating the median, interquartile range and the 95% confidence interval shown by the whiskers. Dots outside the boxplot represent the datapoints that are more than 1.5 times the upper quartile.

### Linkage map construction

To support QTL analysis of the observed differences, we generated SNP genotyping profiles for the Lupa x BGE 016880 population through a RADseq approach and used these to generate a genetic linkage map. A total of 4073 SNP markers were mapped into six linkage groups (LGs; **Fig. 2A and Supplementary file S3**). These groups were designated LG1 through LG6 in accordance with their relationship to chromosomes of the reference genome. Markers from chromosome 7 mapped with those from chromosome 2 in LG2, a pseudo-linkage most likely due to a chromosomal rearrangement in one parent relative to the other. Rearrangements have previously been reported in Lens interspecies crosses (Ladizinsky, 1979) and are also apparent in our recent genome assembly efforts across *Lens* species (Ramsay *et al*., 2019). Respective marker positions between the linkage map and the reference genome are shown is **Fig. 2B**. A heat map of pairwise recombination fractions and LOD scores between all pairs of markers was visualized to check the quality of the linkage map. The tight linkage of the markers within each LG indicated the reliability of the developed linkage map (**Fig 2C**). The total length of the linkage map spanned 5923.3 cM with LG2 being the largest and LG1 the smallest. The number of markers per linkage group varied from 496 in LG1 to 958 in LG2, with an average distance between two adjacent markers of 1.5 cM and the largest gap at 9.4 cM in LG5 (**Supplementary file S1_Table S2**).

**Fig. 2.**
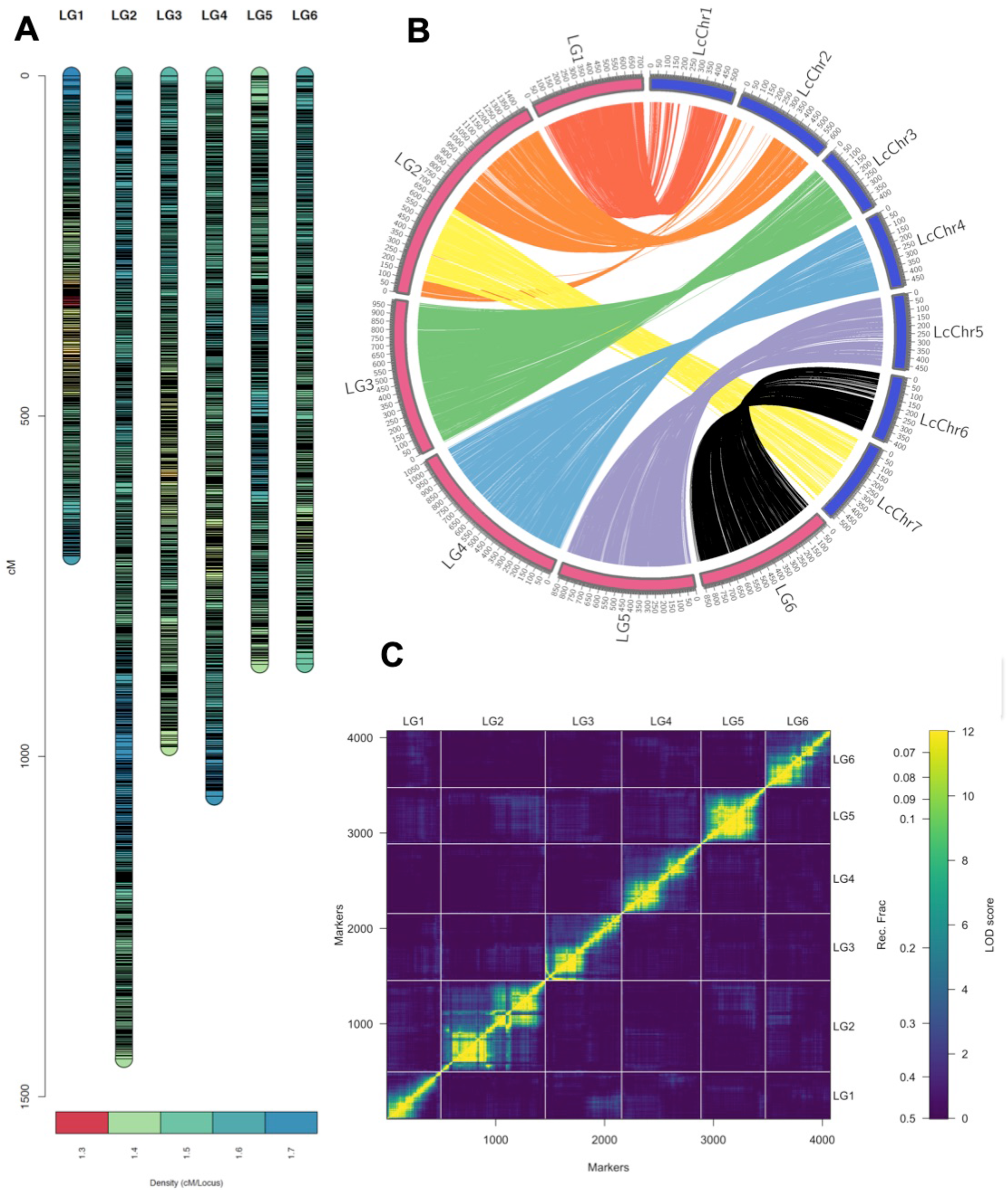
A: Genetic linkage map of a lentil interspecific RIL population developed from a cross between *L. culinaris* cv. Lupa and *L. orientalis* BGE 016880. Scale bar at the left of the linkage map is in centimorgan (cM). Bottom color scheme showed the density of markers with a sliding window of 30 markers for calculation. B: Circular representation of the markers on the genetic linkage map of the RIL population and their respective positions on *Lens culinaris* reference genome. The scale on the outer ring for genetic linkage map (LG1-LG6) is cM and for the reference genome (LcuChr1-LcuChr7) it is million base pairs (Mbp). C: Validation of the map using pairwise linkage information. Plot shows the estimated recombination fractions and LOD scores for all pairs of markers after ordering the markers on each linkage group. The recombination fractions are in the upper left triangle while the LOD scores are in the lower right triangle. Estimates are plotted as a heat map with dark blue signifying no linkage and yellow representing tight linkage with low RF and large LOD. The diagonal yellow line indicates good linkage within each LG. LG2 is the one that is complicated by pseudolinkage of markers from 2 different *L. culinaris* chromosomes.

### Genetic analysis of flowering time and sensitivity to R/FR

We detected six QTLs for flowering time under high R/FR light conditions (*qDTFH-1A, qDTFH-1B, qDTFH-3, qDTFH-5, qDTFH-6A* and *qDTFH-6B*), distributed across four linkage groups (LG 1, 3, 5, and 6) (**Fig. 3**). Among these, one locus (*qDTFH-6A*) explained the majority of the variation (63%), with the other five each contributing less than 10% each (**Table 1**). This QTL showed negative additive effect on days to flowering under high R/FR light quality, which indicated its effect was contributed by the allele from wild parent *L. orientalis* BGE 016880. Under low R/FR light conditions, we detected only four QTLs for flowering time, across LG1 (*qDTFL-1*) and LG6 (*qDTFL-6A, qDTFL-6B, qDTFL-6C*) (**Fig. 3**). As in high R/FR conditions, the overall effect was dominated by one locus (*qDTFL-6A*), which explained 46% of the variance, with the three other loci contributing between 7 and 15% each (**Table 1**). Once again, the wild BGE 016880 allele contributed to the effect at this major locus.

**Fig. 3.**
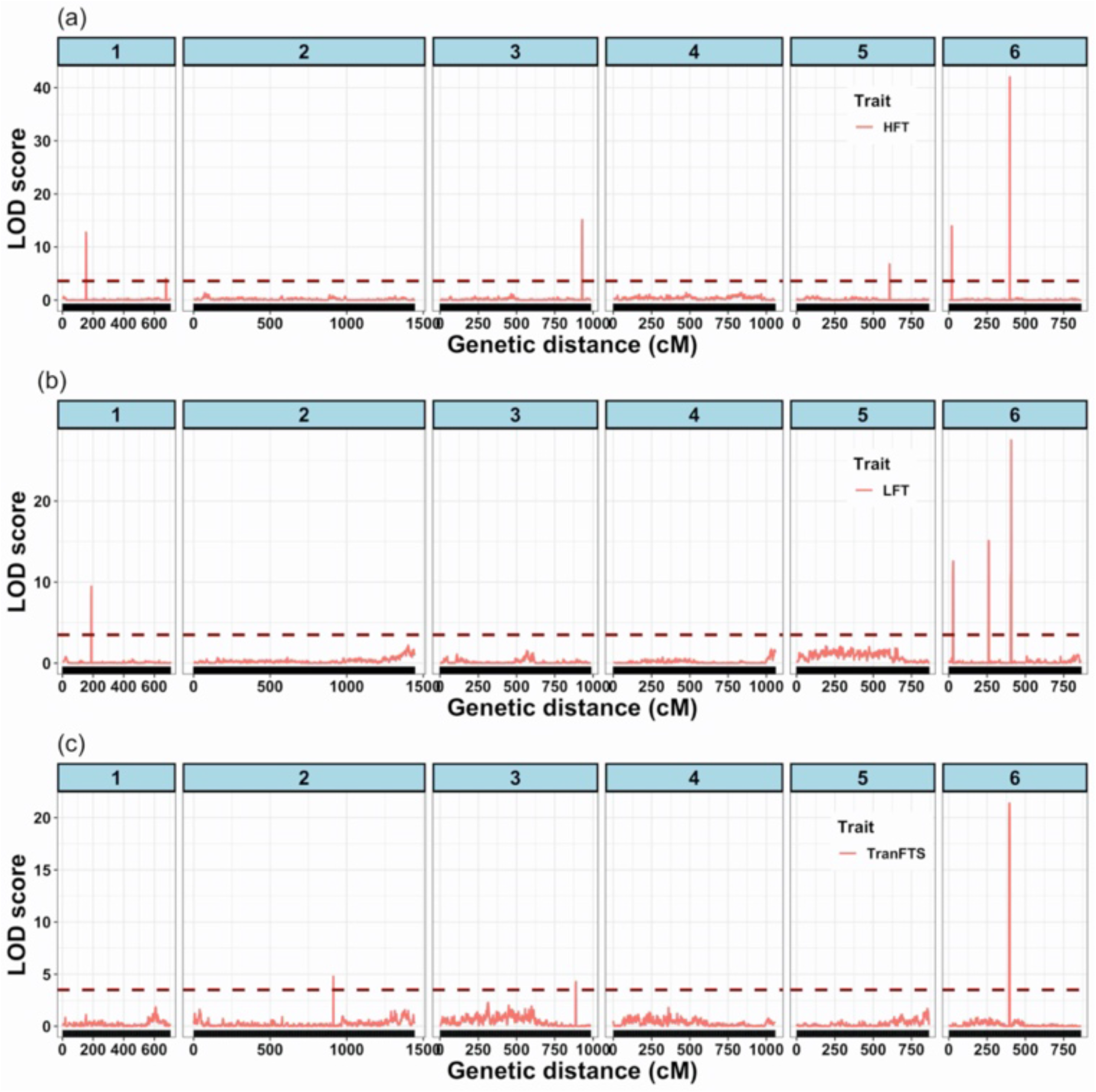
QTLs identified for flower time under two different light quality environments (a – high R/FR and b - low R/FR) and flower time sensitivity to this light quality change (c) in a lentil interspecific RIL population developed from a cross between *L. culinaris* cv. Lupa and *L. orientalis* BGE 016880. The horizontal dashed line on each graph represents the threshold LOD score for QTL identification after 1000 permutation test.

**Table 1.**
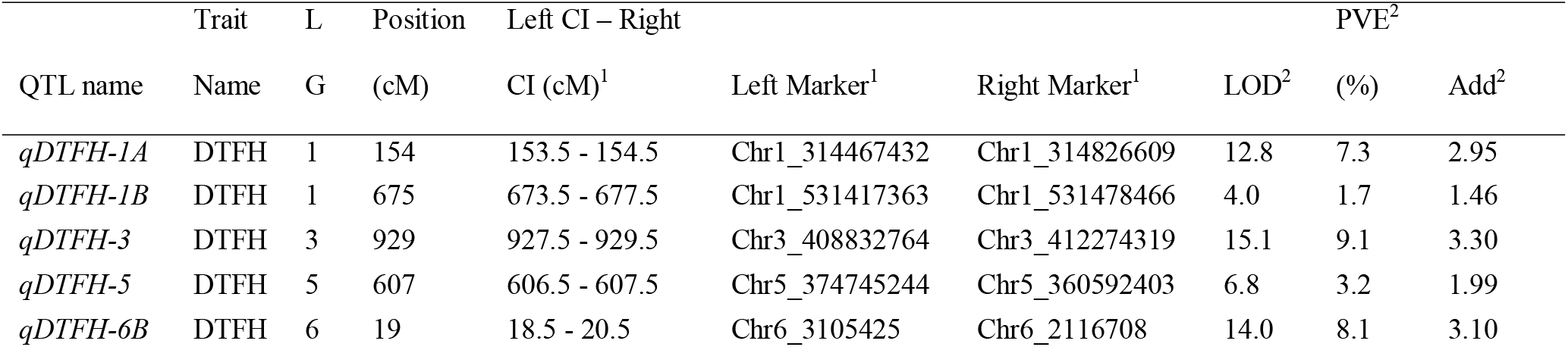
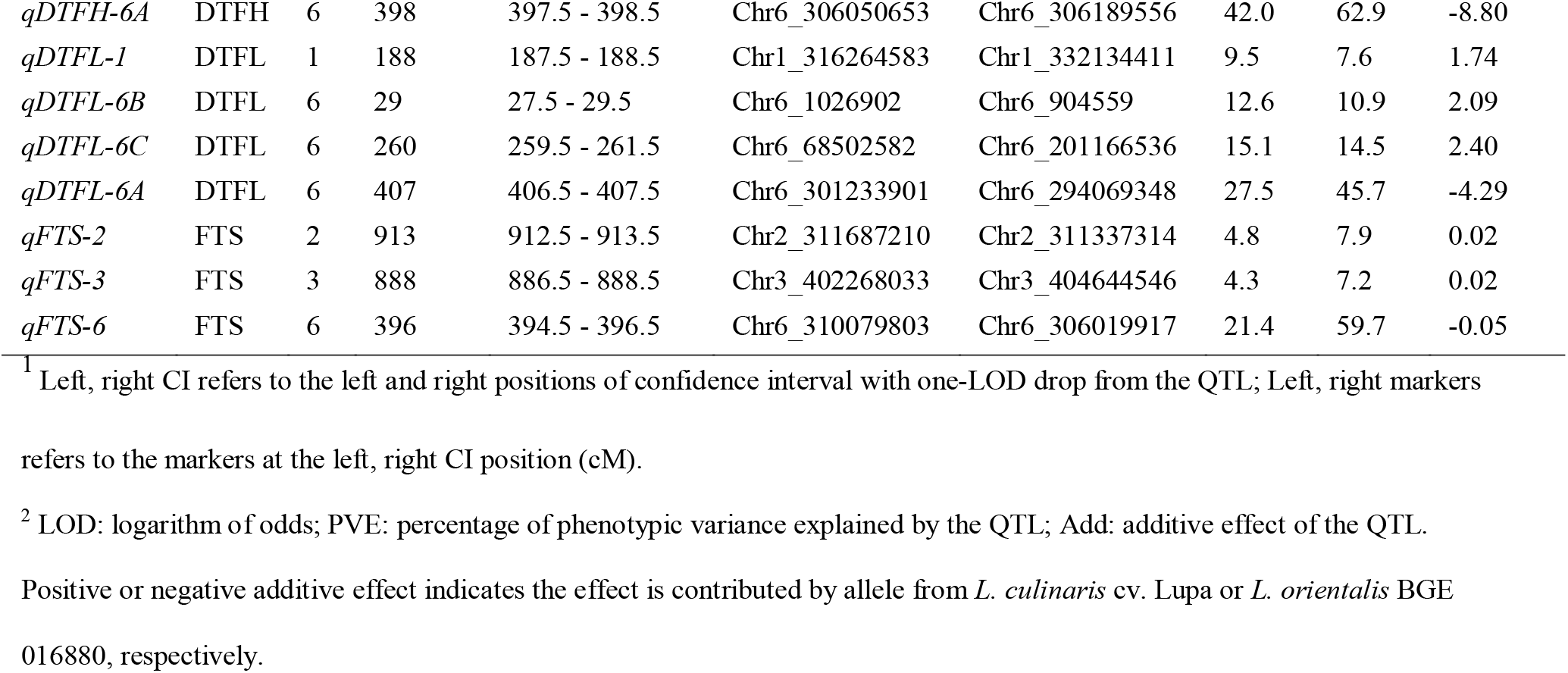
QTLs controlling flowering time under different light quality environments and flower time sensitivity of a lentil interspecific recombinant inbred line population developed from *L. culinaris* cv. Lupa and *L. orientalis* BGE 016880. DTFH - flowering time under high red/far-red light quality environment; DTFL - flowering time under low red/far-red light quality environment, and FTS - flower time sensitivity towards different light quality environments.

Analysis of the flowering time response to R/FR (in terms of the FTS index) revealed three QTLs on LG 2, 3, and 6 (*qFTS-2, qFTS-3, qFTS-6*) (**Fig. 3**). Locus *qFTS-6* was located in a similar region of LG6 to *qDTFH-6A* and *qDTFL-6A* and explained 60% of the phenotypic variance with the contribution from the wild BGE 016880 allele. The other two loci each explained <10% each.

The major locus for each of the three traits (i.e. DTFH, DTFL and FTS) overlapped on LG6 (**Figure 3)**, raising the possibility that these three loci might reflect the contribution of a single underlying gene. Across the three traits, three other pairs of loci showed similar close co-location: on LG1 (*qDTFH1A* and *qDTFL1*), LG3 (*qDTFH-3* and *qFTS-3*) and LG6 (*qDTFH-6B* and *qDTFL-6B*), also suggesting a possible common genetic basis for the loci in each pair. Finally, four of the detected loci (one for each trait) were in unique positions on LG1 (*qDTFH-1B*), LG2 (*qFTS-2*), LG5 (*qDTFH-5*) and LG6 (*qDTFL-6C*). Overall, our analysis across both light quality conditions revealed a minimum of seven distinct QTLs contributing to the variability in DTF under these experimental conditions.

### *De novo* assembly of transcriptome

From what we have done so far based on the genome assemblies of different *Lens* species and intensive genotyping of intra-, inter-specific lentil RIL populations, we know that genome size difference exists among *Lens* species and chromosomal rearrangements are common even within cultivated lentil. BGE 016880 is an accession of *L. orientalis* while Lupa is a Spanish cultivar (Fratini *et al*., 2007). To remove bias that could result from using the *L. culinaris* cv. CDC Redberry reference genome for differential gene expression analysis, we carried out *de novo* assembly of the transcriptomes from RNAseq data for both BGE 016880 and Lupa.

A total of 705.1 MB paired-reads of the transcriptomic data were generated using the Illumina HiSeq 2500 platform (paired end of 125bp*2) of which 359.4 MB belonged to *L. orientalis* BGE 016880 and 345.7 MB belonged to *L. culinaris* cv. Lupa (**Supplementary file S4**). After quality filtering and pre-processing of reads, 97.8% of the reads for both *L. orientalis* BGE 016880 and *L. culinaris* cv. Lupa were kept for further analysis. These high-quality reads of 30 samples from five developmental stages were pooled together for *L. orientalis* BGE 016880 and *L. culinaris* cv. Lupa, respectively, and used for *de novo* transcriptome assembly. Trinity assembler generated 138108 transcripts (78707 raw genes) with N50 of 2255bp for *L. orientalis* BGE 016880, and 135723 transcripts (76595 raw genes) with N50 of 2252bp for *L. culinaris* cv. Lupa. (**Supplementary file S1_Table S3**).

### Differential gene expression analysis

*De novo* assembled transcriptomes of BGE 016880 and Lupa were used to quantify the abundance of the transcripts for all samples from BGE 016880 and Lupa respectively using Salmon package (Patro *et al*., 2017). 3D RNA-seq pipeline (Calixto *et al*., 2018; Guo *et al*., 2019) was then used for the analysis of differential gene expression and a detailed method can be found in **Supplementary file S1_Supplementary_Methods**.

A total of 2573 differentially expressed genes (DEGs) and 1367 DEGs were obtained from the contrast sets between the low R/FR light environment and high R/FR environment for five different growth stages of *L. orientalis* BGE 016880 and *L. culinaris* cv. Lupa (**Supplementary file S5**). Out of the 2573 DEGs from BGE 016880, 933, 1061, 64, 139 and 376 DEGs were observed in T1, T2, T3, T4, and T5 stage, respectively (**Supplementary file S1_Figure S3A**). Out of the 1367 DEGs from Lupa, 434, 84, 232, 362 and 255 DEGs were observed in these five stages, respectively (**Supplementary file S1_Figure S3B**). Up and down-regulated genes at these five stages were quite different for both *L. orientalis* BGE 016880 and *L. culinaris* cv. Lupa (**Fig. 4**), however most DEGs were identified about 2 weeks before flowering which corresponds to the transition from vegetative growth to reproductive growth for both BGE 016880 and Lupa. When samples from the five development stages were pooled together to look at the DEGs between low R/FR and high R/FR light quality, 297, and 156 were identified from *L. orientalis* BGE 016880 and *L. culinaris* cv. Lupa, respectively (**Supplementary file S5**). Differentially expressed genes in *L. orientalis* BGE 016880 were almost double compared to those in *L. culinaris* cv. Lupa between low R/FR and high R/FR light quality environments.

**Fig. 4.**
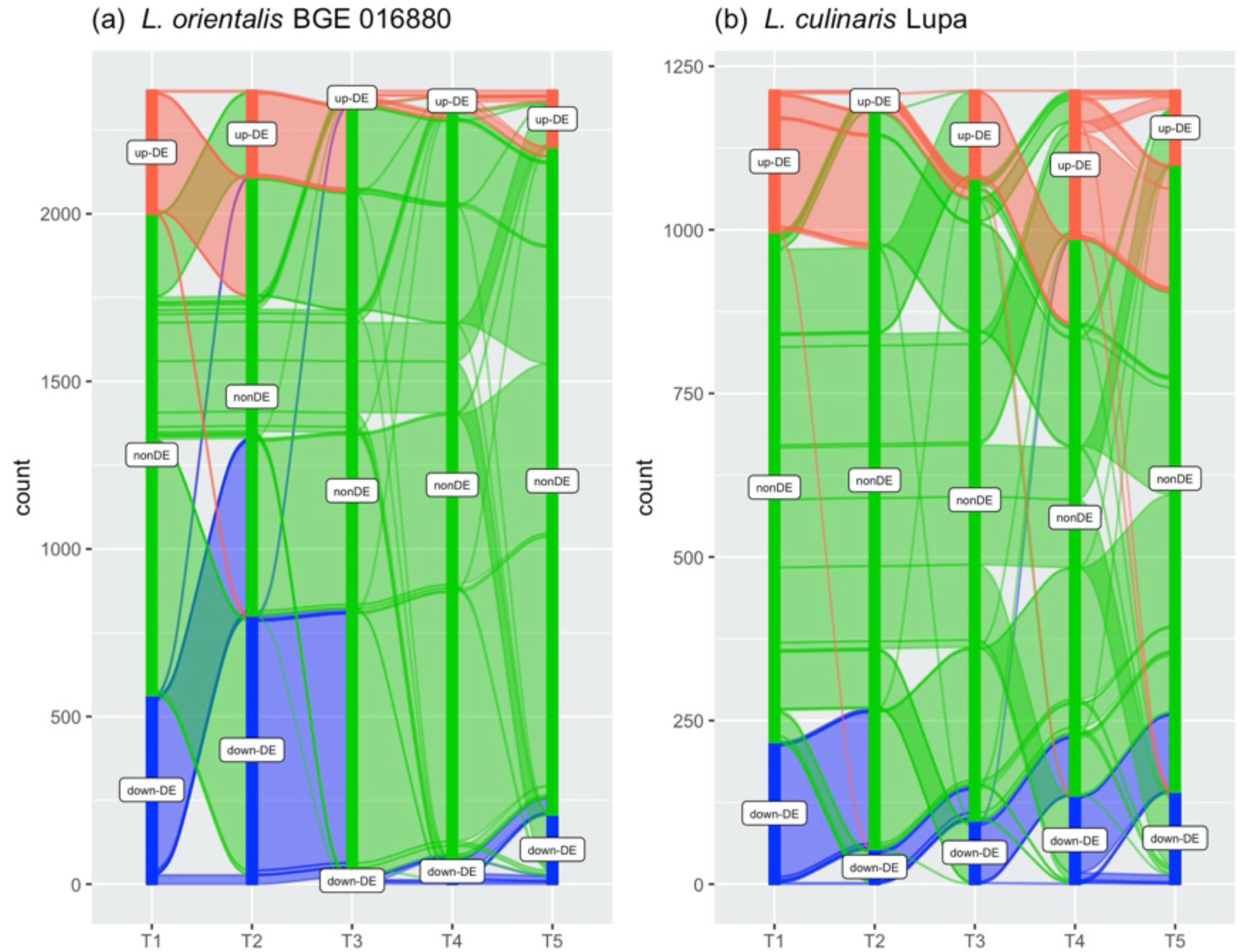
Alluvial plots showing up-regulated genes (red flow), non-differentially expressed genes (green flow) and down-regulated genes (blue flow) from the contrast between low R/FR and high R/FR from five different growth stages for *L. orientalis* BGE016880 (a) and *L. culinaris* cv. Lupa (b).

### Identification of flowering DEGs

We further examined the DEGs for possible functional links to flowering time regulation. To do this, we used the list of 244 lentil genes that we curated based on the reference genome (**Supplementary file S2**) and examined their representation within the DEG set using a custom Perl script (**Supplementary file S6**). This analysis identified a total of 82 genes as candidate flowering-related genes among the DEG sets identified for *L. orientalis* BGE 016880 and 62 from the *L. culinaris* cv. Lupa DEG sets. The number of flowering-related DEGs associated with each of the five development stages were as follows: T1-15, T2 - 24, T3 - 13, T4 - 12 and T5 - 18 for BGE 016880. When sorting flowering-related DEGs within the development stages for Lupa, we saw the following numbers: T1 - 10, T2 - 4, T3 - 15, T4 - 17 and T5 - 16. Flowering-related DEGs again showed stage specific regulation in both BGE 016880 and Lupa.

Since the flowering times for both BGE 016880 and Lupa were significantly different and spread from 27 to 60 days after emergence (**Supplementary file S1_Fig. S1**), we decided to look at flower DEGs from the DEG set of samples from five development stages, instead of any single stage, to understand the flower time regulation between low R/FR and high R/FR light quality environments. 17 flower DEGs were identified in both BGE 016880 and Lupa, out of 297 and 156 DEGs respectively. Of these, 12 were common to both BGE 016880 and Lupa, while five others were unique to DEG set for one or the other line (**Table 2**).

**Table 2.**
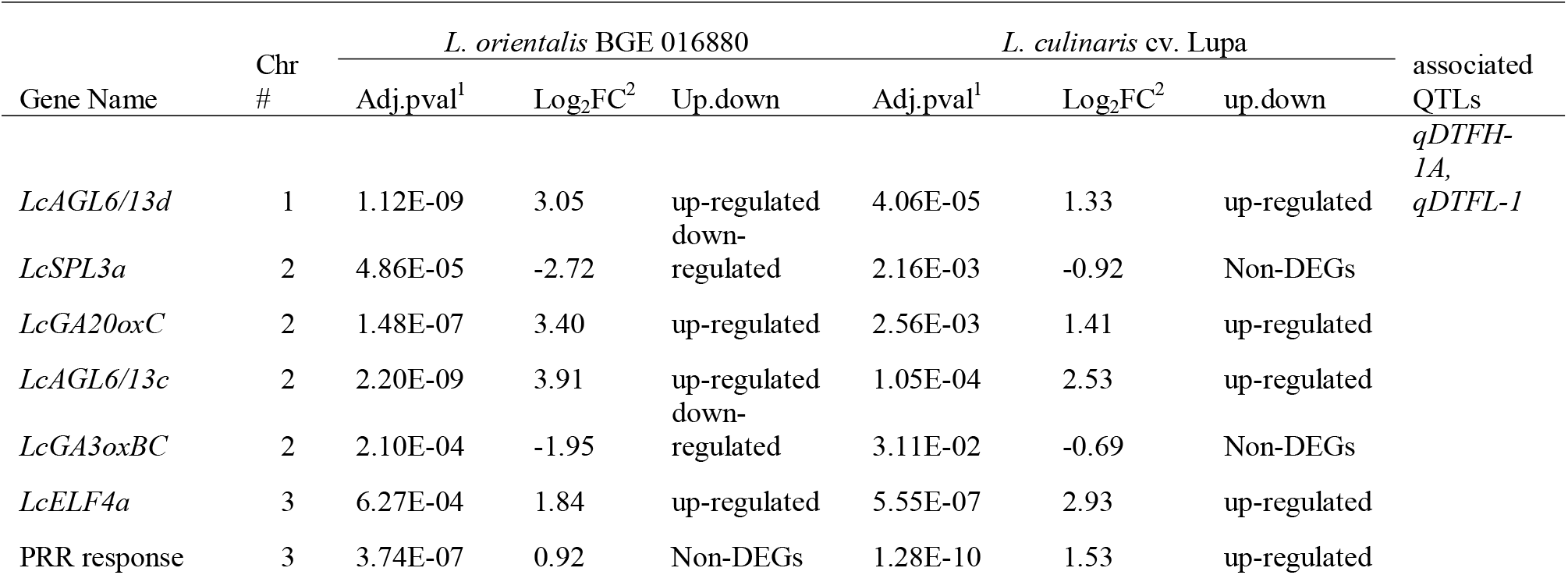
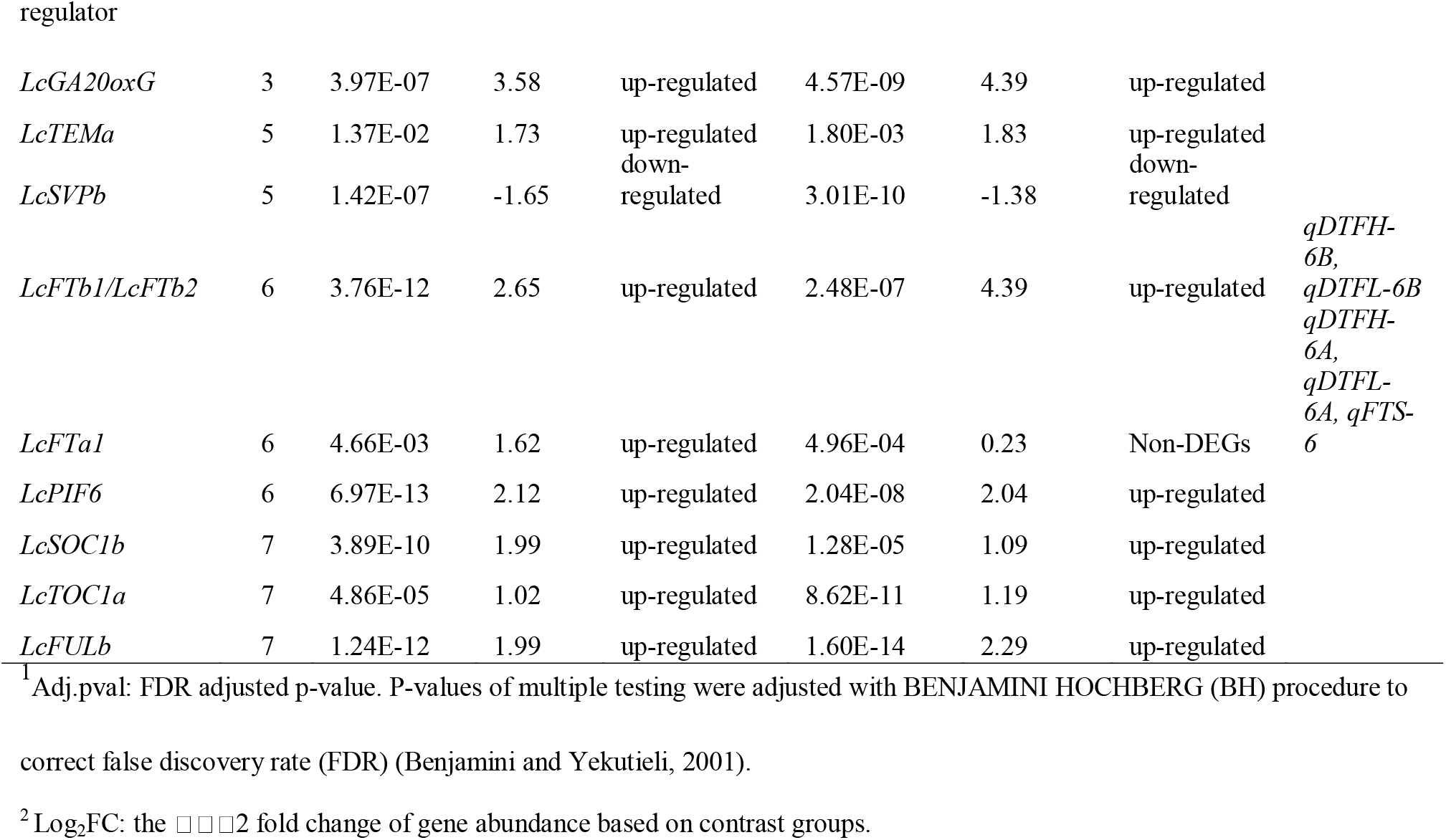
Flowering DEGs from the contrast set between low red/far-red ratio and high red/far-red ratio for both *L. orientalis* BGE016880 and *L. culinaris* cv. Lupa and possible related QTLs identified from a lentil interspecific RIL population developed from *L. culinaris* cv. Lupa and *L. orientalis* BGE 016880. A gene was significantly differentially expressed if it had adjusted p-value < 0.05 and |Log_2_FC| ≥ 1.

### Verification of flowering DEGs through RT-qPCR

We next validated results from RNAseq analysis by RT-qPCR, for selected genes including Flowering DEGs in the QTL regions, non-DEG flowering genes within the QTL regions, and flowering DEGs that were not within QTL regions. From the subsets of flowering-related DEGs, we selected five genes that were located within or nearby the QTL confidence intervals (**Table 2**). The major loci *qDTFH-6A, qDTFL-6A* and *qFTS-6* in the central region of LG6 co-located with three *FT* homologs: *LcFTa1, LcFTa2* and *LcFTc* (Lcu.2RBY.6g043850, Lcu.2RBY.6g043870, Lcu.2RBY.6g043940), and a light-regulated transducin/WD-like repeat-protein (LWD) gene ortholog *LcLWD1* (Lcu.2RBY.6g043520). Of these, *LcFTa1* was the only one represented in the DEG set in the wild parent, but not in the cultivated parent. Another cluster of *FT* homologs *LcFTb1* and *LcFTb2* (Lcu.2RBY.6g000730 and Lcu.2RBY.6g000760) were located near the *qDTFH-6B* and *qDTFL-6B* loci at the top of LG6. A MADS-box gene in the *AGAMOUS-LIKE 6* (AGL6) clade (Lcu.2RBY.1g040350/ *LcAGL6/13d*) was the closest flowering-related DEG to the loci *qDTFH-1A* and *qDTFL-1*. We also selected one gene *LcELF4a* (Lcu.2RBY.3g037650), a flowering-related DEG not located within or nearby any QTL region.

The results were consistent with, and further supported our RNAseq results (**Fig. 5**). For the major QTL region in the middle of LG6, we confirmed the strong upregulation of *LcFTa1* in response to low R/FR in *L. orientalis* BGE 016880 but not in *L. culinaris* cv. Lupa. In the same region, *LcLWD1* and Lc*FTa2* had lower (or no) expression in *L. orientalis* BGE 016880, but similar expression level in *L. culinaris* cv. Lupa under different light quality environments, whereas *LcFTc* expression was unaffected by R/FR (**Fig. 5**). For the QTL region at the top of LG6, the DEGs *FTb1* and *FTb2* showed consistent increased expression under low R/FR light quality in both *L. orientalis* BGE 016880 and *L. culinaris* cv. Lupa. *LcELF4a* was not within or nearby any QTL region, however, it was identified as a flowering DEG and RT-qPCR result also supported this. This could mean that additional loci with small effects were not identified, a common limitation of QTL mapping (Myles and Wayne, 2008).

**Fig. 5.**
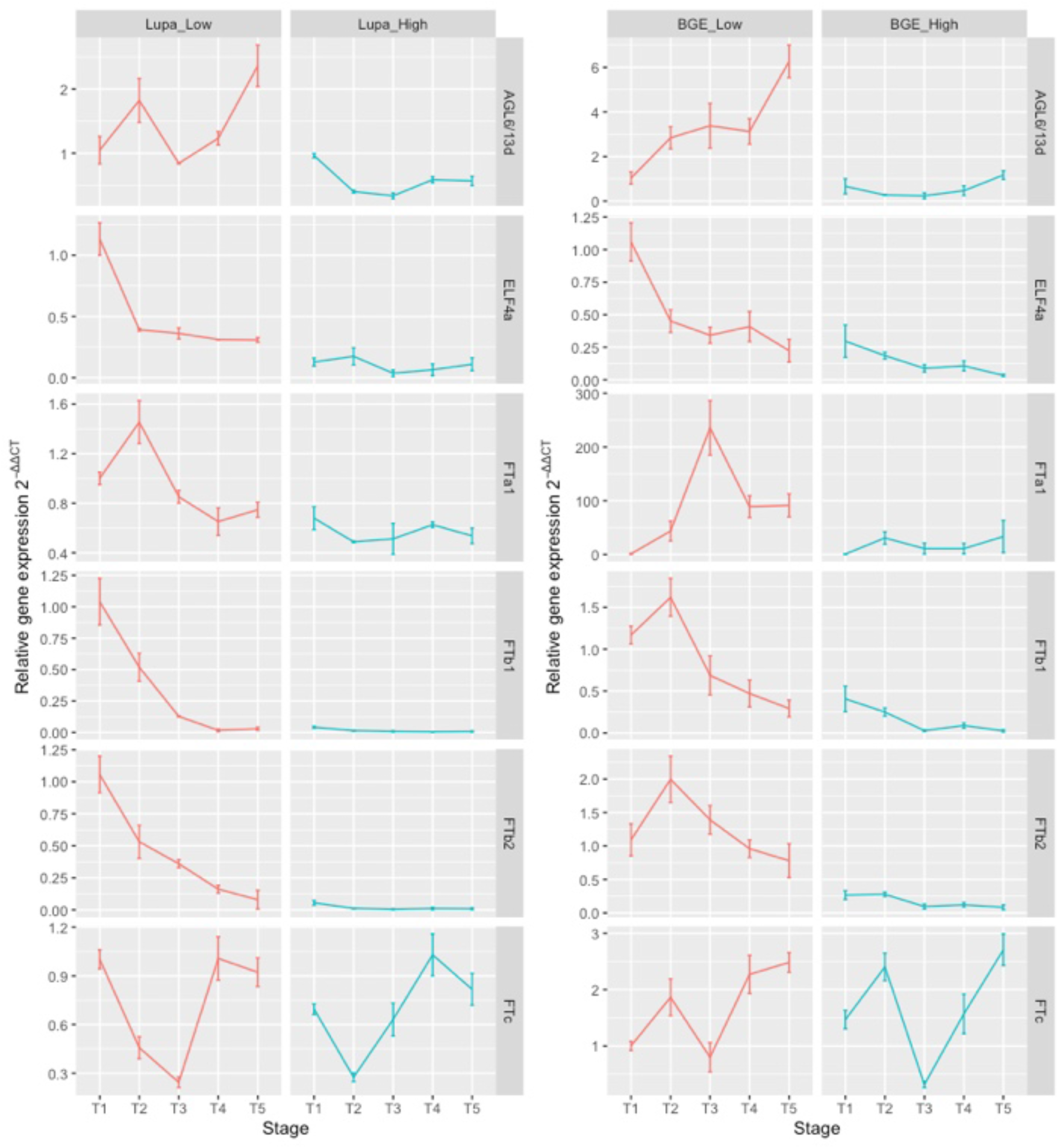
Relative expression of flowering-related genes from the samples grown at low red/far-red ratio and high red/far-res ratio from five different growth stages for both *L. orientalis* BGE016880 and *L. culinaris* cv. Lupa assessed using RT-qPCR. Sample collection started 2 weeks after emergence and continued once a week for 5 weeks. Samples were taken at the same time of the day for each collection. Actin was used as the reference gene for the normalization of the data and the delta-delta CT method was adopted to derive the relative gene expression value 2^-ΔΔCT^ using samples from T1 stage at low R/FR light quality environment as reference samples. Values represent the mean of biological replicates with their corresponding standard deviation.

## DISCUSSION

Optimization of seed yield and reproductive success in nature and in agriculture depends to a large extent on the appropriate seasonal control of flowering. (Jung and Müller, 2009). Light is an important daily, seasonal and spatial variable for plants, and flowering time is not only affected by light duration (photoperiod), but also by light quality (Adams *et al*., 2009; Casal, 2013). In particular a reduction in R/FR occurs in light reflected from green plant tissue, provides cues about prospective competition from neighbouring plants, and leads to a set of morphological responses including an acceleration of flowering (Whitelam and Devlin, 1997). We have previously reported species and genotype differences in response to light quality changes within the genus *Lens*, where wild lentils seemed to be less sensitive than cultivated types and therefore may perform better under variable light quality (Yuan *et al*., 2017).

The parents of the interspecific RIL population used in this study have contrasting flowering responses to changes in light quality, with the wild parent, BGE 016880, being less sensitive to changes than is the cultivated parent, Lupa. As expected, the RILs within the population exhibited segregation for flowering time under different light quality environments, as well as flowering time sensitivity towards different light qualities. The presence of less sensitive RILs than the wild parent BGE 016880, and more sensitive RILs than the cultivated parent, suggests that alleles from both parents have contributed to the response and rearrangement of alleles lead to transgressive segregation in the RIL population.

In our current study, 4073 SNP markers were distributed across 6 linkage groups with an average marker density of 1.5 cM. The physical size of the lentil genome is estimated to be 4086 Mb (Arumuganathan and Earle, 1991). Based on the distance of the map developed in this study, an average of 0.69 Mb/cM was covered. Some similar linkage maps generated in lentil RIL populations had average coverages of 5.9 Mb/cM (Fedoruk *et al*., 2013), 2.13-3.14 Mb/cM (Sudheesh *et al*., 2016) or 1.01 Mb/cM (Ates *et al*., 2016) using SNP markers generated from other approaches. It appears that the amount of RADseq-generated SNP markers used in the current map may help to improve coverage of the lentil genome.

There are few reports on flowering time related QTL studies in lentil, and most have used different marker systems, making it difficult to find consensus across studies (Sarker *et al*., 1999; Fratini *et al*., 2007; Tullu *et al*., 2008; Fedoruk *et al*., 2013). So far, there have been no reported studies on the genetics behind flowering time and light quality change in lentil, although the identity of the major Sn locus as an ortholog of the circadian clock gene ELF3a (Sarker *et al*., 1999; Weller *et al*., 2012) suggests a possible influence on both photoperiod and light quality sensitivity (Jiménez-Gómez *et al*., 2010; Weller *et al*., 2012). In our current study, thirteen QTLs were identified in the RIL population for days to flowering under high R/FR light quality, days to flowering under low R/FR light quality, and flowering time sensitivity towards light quality change.

Studies using comparative genetics as well as functional genomics have shown that most major flowering time genes are well conserved between Arabidopsis and a large range of crop species including legumes (Hecht *et al*., 2005; Roux *et al*., 2006; Kim *et al*., 2013; Weller and Ortega, 2015). Using in-house curated 244 lentil homologs of *Arabidopsis thaliana* and *Medicago truncatula* flowering related genes, we were able to look at the corresponding QTL regions in the lentil reference genome and associated them with flower DEGs. Three major QTLs identified in the study were within a similar region in the middle of LG6, where a tandem array of florigen (FT) genes: *LcFTa1, LcFTa2* and *LcFTc*, as well as a light-regulated transducin/WD-like repeat-protein (*LcLWD1*), are located, although only one (*LcFTa1)* showed significant differential expression within the region. These three QTLs explained more than 45% phenotypic variance respectively and were in a region syntenic with a section of Medicago chromosome 7 and pea linkage group V, containing a similar tandem array of *FTa1, FTa2* and *FTc* genes (Hecht et al. 2005, Weller and Ortega, 2015). This region is related to control of flowering time in several temperate legumes, including pea (Hecht *et al*., 2011), *M. truncatula* (Laurie *et al*., 2011), faba bean (Cruz-Izquierdo *et al*., 2012), narrow-leafed lupin (Nelson *et al*., 2017), and chickpea (Ortega *et al*., 2019). When *de novo* assembled transcriptomes of *L. orientalis* BGE016880 and *L. culinaris* cv. Lupa were aligned with our reference genome, the *LcFTa2* transcript could not be detected in *L. orientalis* BGE016880, implying a possible deletion, and suggesting that this gene is not likely to be involved in the response to changes in light quality in this interspecific RIL population. Interestingly, a similar deletion was also observed to associate with a QTL conferring early flowering in chickpea (Ortega *et al*., 2019), but was also not considered likely to explain the effect, in view of the generally promotive effects of *FT* genes on flowering (Laurie *et al*., 2011; Hecht *et al*., 2011).

Another cluster of QTLs in a distinct region at the top of LG6 was shown to co-locate with two other differentially expressed flowering genes, the florigen genes *LcFTb1* and *LcFTb2*. In both pea and Medicago, orthologs of these genes have also been implicated in flowering time control through genetic and gene expression studies (Laurie *et al*., 2011; Hecht *et al*., 2011; Ridge *et al*., 2017; Putterill *et al*., 2019; Jaudal *et al*., 2020). As a flowering pathway integrator, Flowering Locus T genes promote the transition to flowering and play a key role in plant adaptation and crop improvement (Wickland and Hanzawa, 2015). In the related temperate legume subterranean clover, *FTb2* has also been implicated in the acceleration of flowering by low R/FR through gene expression studies (Pazos-Navarro *et al*., 2018).

Two other QTLs identified were within the same region on LG1 where Agamous-like MADS-box protein AGL6/13d (*LcAGL6/13d*) was the sole flowering-related DEG. AGL6 represents a subfamily of MADS-box transcription factor genes and have various roles regarding flowering time and flower development (Dreni and Zhang, 2016). In Arabidopsis, the activation of *AGL6* is associated with the down-regulation of floral repressor *FLC/MAF* genes, and up-regulation of the floral promoter *FT* (Yoo *et al*., 2011). RT-qPCR analysis confirmed that *LcAGL6/13d* consistently showed high expression under low R/FR light quality compared to high R/FR light quality in both *L. orientalis* BGE016880 and *L. culinaris* cv. Lupa.

FT regulators like Phytochrome Interacting Factors (PIFs) and Short Vegetative Phase (SVP) target FT promoters and non-coding regions. These regulators were also differentially expressed in our study. PIFs belong to a class of basic helix-loop-helix (bHLH) transcription factors and are involved in promoting the transition to flowering by acting upstream of FT and *TWIN SISTER of FT* (*TSF*) under low R/FR in Arabidopsis (Galvao *et al*., 2019). In soybean, *PIF3* has been identified as a shade-responsive gene through an RNAseq experiment (Horvath *et al*., 2015). *SVP*, or Agamous-like MADS-box protein AGL22 is a negative regulator of flowering in Arabidopsis (Hartmann *et al*., 2000) and can bind to the promoters of *SOC1* and *FT* for transcriptional repression (Li *et al*., 2008). *SVP* expression decreased after a Gibberellin (GA) treatment, which suggested the involvement of GA on flowering could be partly mediated through *SVP* (Li *et al*., 2008). Two other members of MADS-box transcription factor genes, Suppressor of overexpression of constans 1 (*SOC1*) and FRUITFULL (*FUL*), were also identified as flowering-related DEGs in our study. As a floral pathway integrator, *SOC1* (Agamous-like MADS-box protein AGL20) incorporates multiple flowering signals to promote flowering and can be induced by GA, while repressed by *FLC* and *SVP* in Arabidopsis (Lee and Lee, 2010). *SOC1* works downstream of *FT* to promote flowering through up-regulation of the gene *LEAFY*, which links floral induction and floral development. A recent study in *Medicago truncatula* showed that *MtSOC1* genes can be induced by *MtFTa1* gene and function redundantly to accelerate flowering (Fudge *et al*., 2018). *FUL*, or Agamous-like MADS-box protein AGL8 has been shown to work together with *SOC1*, while act against the effects of *FLC* and *SVP* to promote flowering (Balanzà *et al*., 2014). The subset of flowering-related genes differentially expressed under low compared to high R/FR light quality was very similar in both *L. orientalis* BGE016880 and *L. culinaris* cv. Lupa, indicating a high similarity in the underlying regulatory processes. However, they differed to some extent both in the level of the expression differences and the stage at which specific differences were detected.

Overall, our results from QTL analysis and gene expression point most clearly to *FTa1* as a probable basis for the observed differences in flowering sensitivity to light quality between *L. orientalis* BGE016880 and *L. culinaris* cv. Lupa. Similar parallel evidence suggests a weaker role for *FTb1/2* genes, and identifies an AGL6/13-like MADS-box transcription factors as a potential candidate for a third QTL. In addition, the wider gene expression dataset suggests a number of other genes including MADS box genes *LcSVPb, LcSOC1b* and *LcFULb*, the bHLH transcription factor *LcPIF6*, and genes related to gibberellin synthesis *LcGA20oxC,G*, may be involved in the light quality response in leaf tissue, either upstream or downstream of *FT* genes (**Fig. 6**). An important characteristic of the flowering regulatory network is that all signaling pathways respond to different endogenous and environmental signals but work together through a delicate and complicated balance to adjust the switch to flower in a changing environment. Our results suggested that *FT* genes together with various MADS-box and bHLH transcription factors played an important role in flowering time regulation under different light quality environments. The identified QTLs and candidate genes provide a foundation for better adaptation to variable light environments in crop improvement of lentil.

**Fig. 6.**
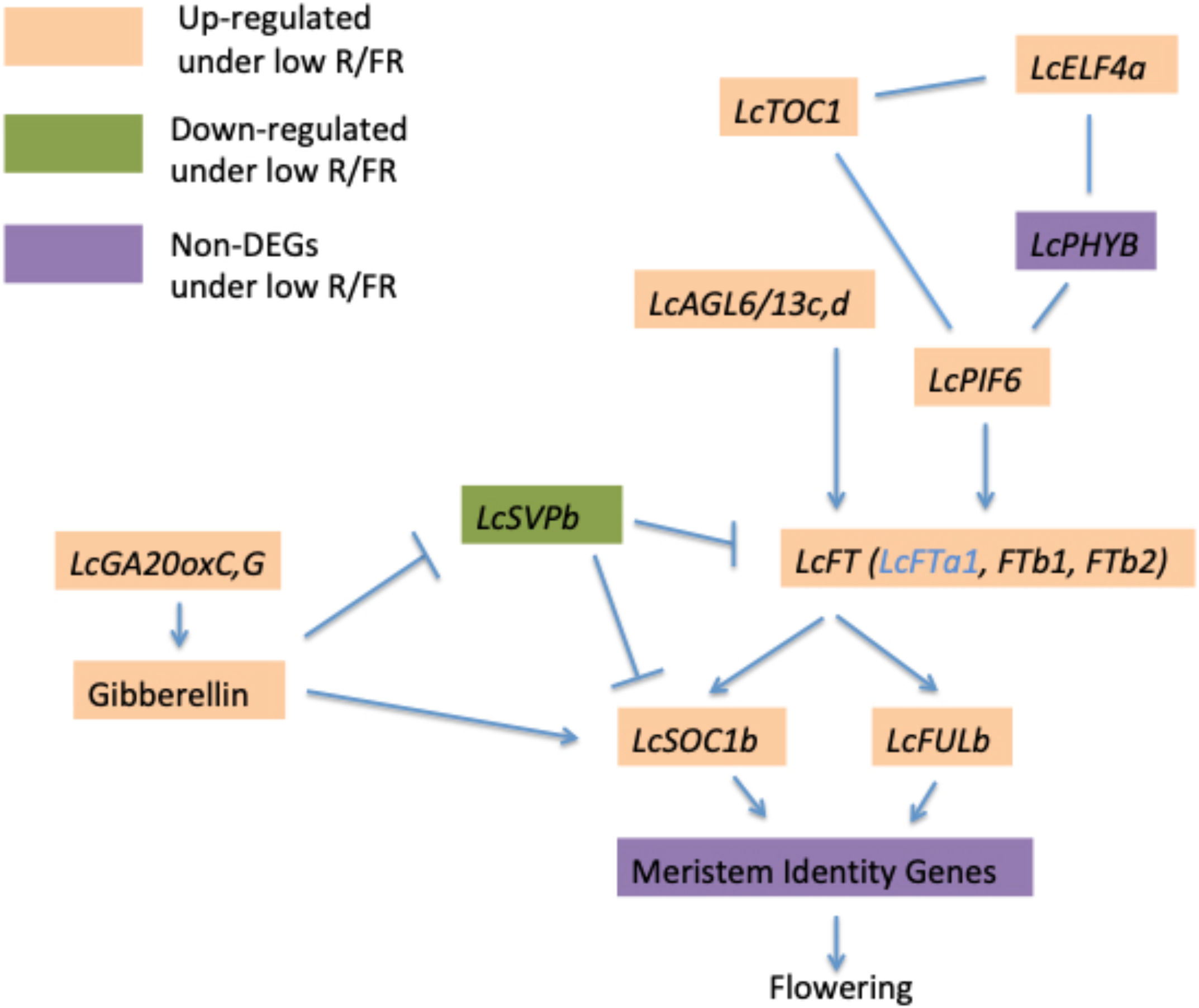
A model for the role and interactions of lentil flower genes under low R/FR light quality environment. This model summarizes the major results from this study and the hypothetical interactions are based on previous network studies on Arabidopsis and other legumes (Putterill *et al*., 2013; BlüMel *et al*., 2015; Weller and Ortega, 2015; Chen *et al*., 2018). Arrows indicate a promoting interaction, a T-end indicates an inhibiting interaction, and a straight line marks an interaction with no firm direction. Gene name shown in blue (*LcFTa1*) was a DEG only in *L. orientalis* BGE 016880.

## ACKNOWLEDGEMENTS

We acknowledge Ms. Devini De Silva, Ms. Akiko Tomita, Ms. Brianna Jansen and Mr. Ricco Tindjau for assistance with the technical work. Technical expertise of the Phytotron staff in the College of Agriculture and Bioresources at the University of Saskatchewan is greatly appreciated.

## FUNDING

The research was conducted as part of the AGILE project (Application of Genomics to Innovation in the Lentil Economy), a Genome Prairie managed project funded by Genome Canada, Western Grains Research Foundation, the Province of Saskatchewan, and the Saskatchewan Pulse Growers.

## Notes

### Competing Interest Statement

The authors have declared no competing interest.

