## Supplementry File 1 for "Genetic and gene expression analysis of flowering time regulation by light quality in lentil"

|  | Days to flower under low R/FR | Days to flower under high R/FR | Difference in Days to Flower between two conditions | Flowering time sensitivity^a^ |
| --- | --- | --- | --- | --- |
| *L.culinaris* cv. Lupa | 36 | 60 | 24 | 0.254 |
| *L.orientalis* BGE 016880 | 27 | 38 | 11 | 0.178 |
| RIL population |  |  |  |  |
| Minimum | 27 | 32 | 5 | 0.074 |
| Maximum | 50 | 70 | 36 | 0.374 |
| Mean | 32 | 51 | 19 | 0.219 |
| Median | 31 | 52 | 20 | 0.222 |

Supplementary Figures

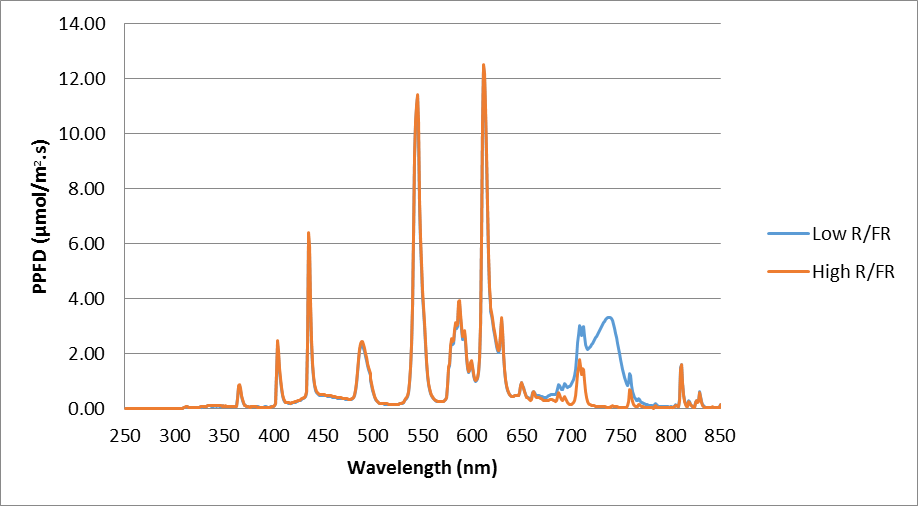

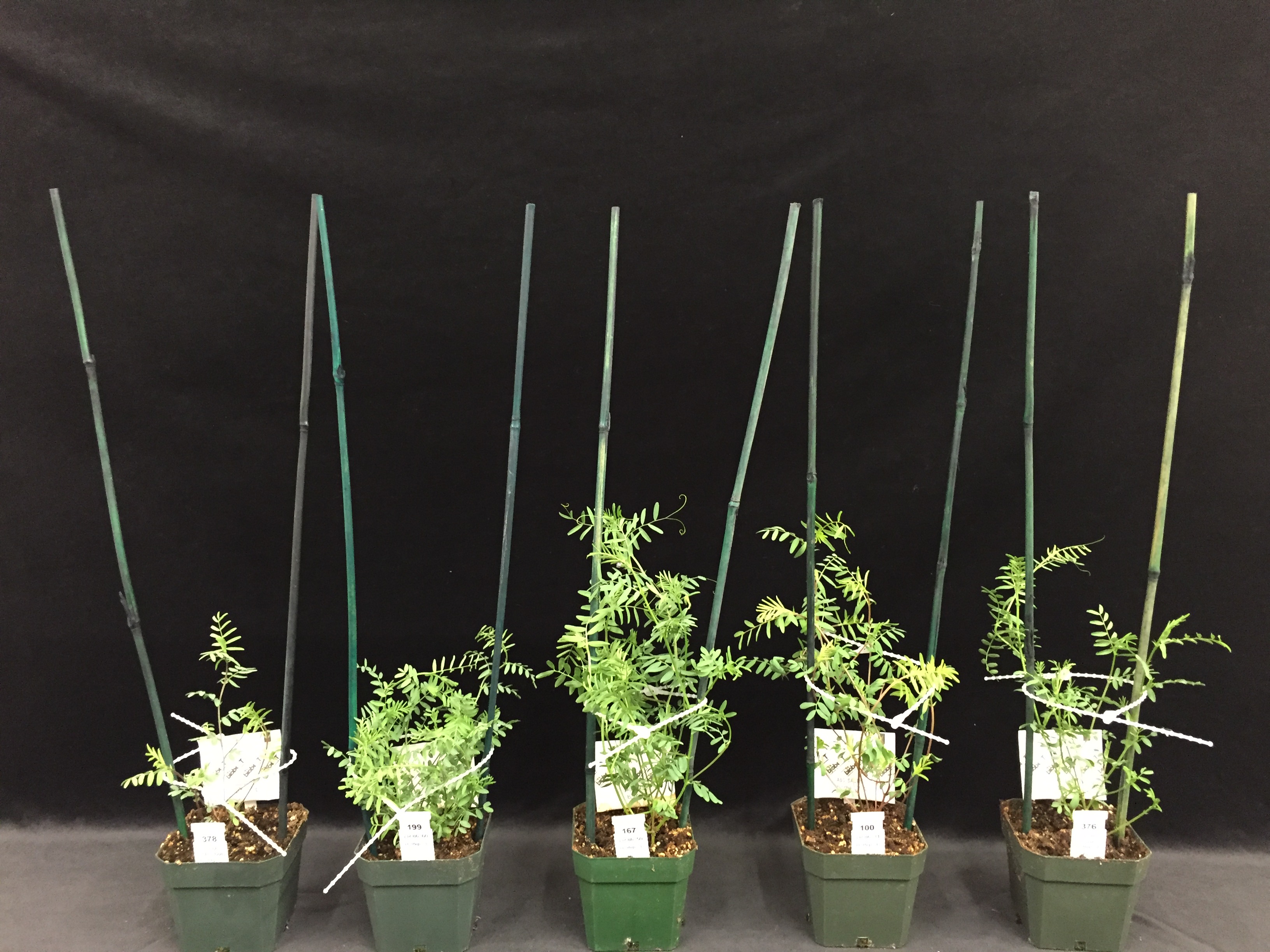

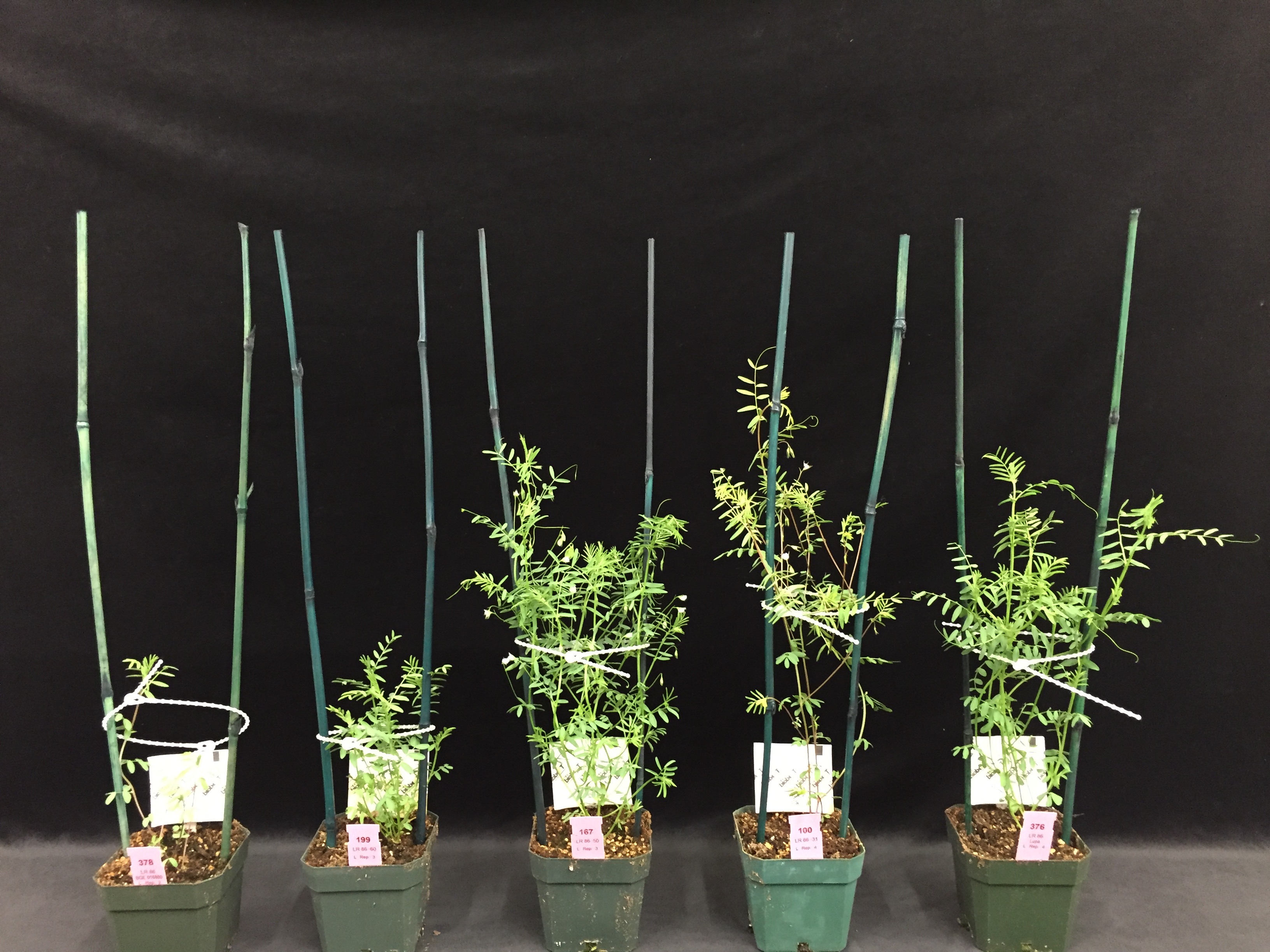

**A**

**C**

**B**

Fig. S1. Spectral distribution of the light quality treatments used to compare the effect of light quality on days to flower of a lentil interspecific recombinant inbred line (RIL) population developed from *L. culinaris* cv. Lupa and *L. orientalis* BGE 016880 (A) and various responses of sub-lines from this interspecific RIL population to light environments differing in red to far-red ratio (R/FR). B: High R/FR light environment. C: Low R/FR light environment. Left to right (in both conditions): Parent *L. or.* BGE 016880, inbred line (IL) 60, IL 50, IL 31, and Parent *L. cu.* Lupa. Photos were taken at 5 weeks after seeding when IL 50 and IL 31 had flowered under low R/FR condition. ^a^ Flowering Time Sensitivity (FTS) was calculated as the ratio of the difference in DTF under the two conditions divided by the sum of the DTF under the two conditions so as to avoid bias due to the underlying differences in DTF. FTS = (DTF_high_-DTF_low_)/(DTF_high_+DTF_low_)

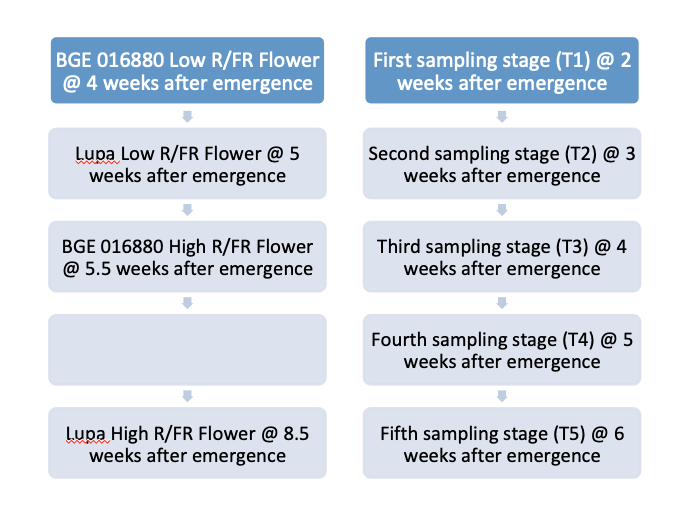

Fig. S2. Relative flowering times for both *L. culinaris* cv. Lupa and *L. orientalis* BGE 016880 under different R/FR light environments and sampling stages used for the RNAseq study.

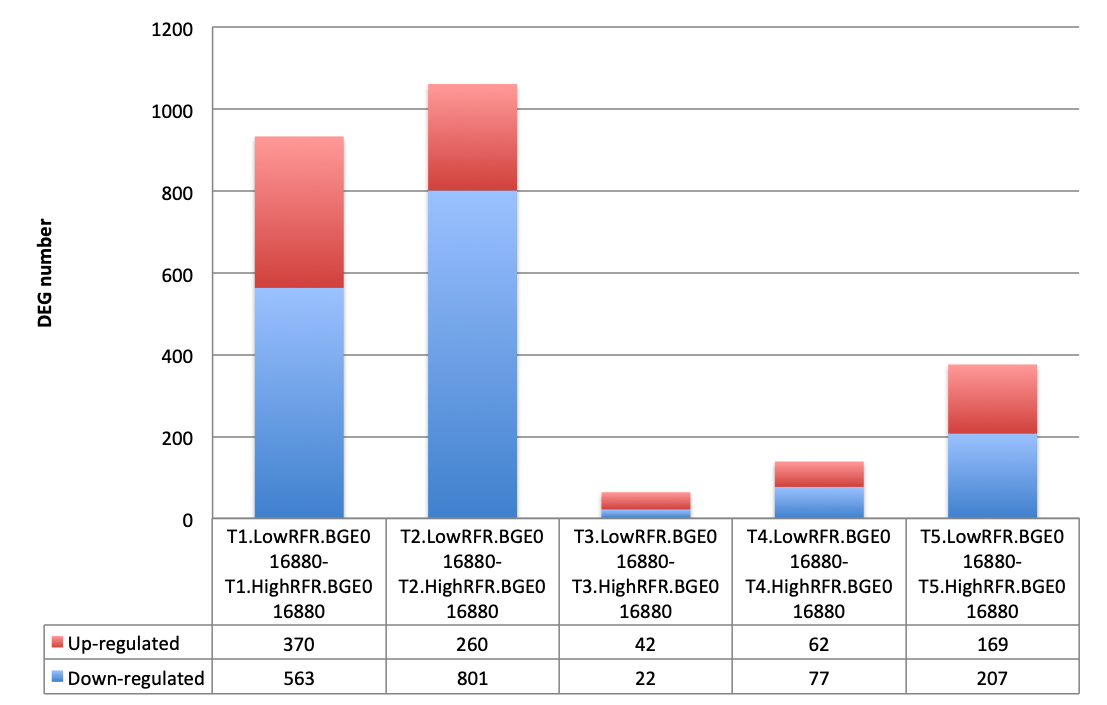

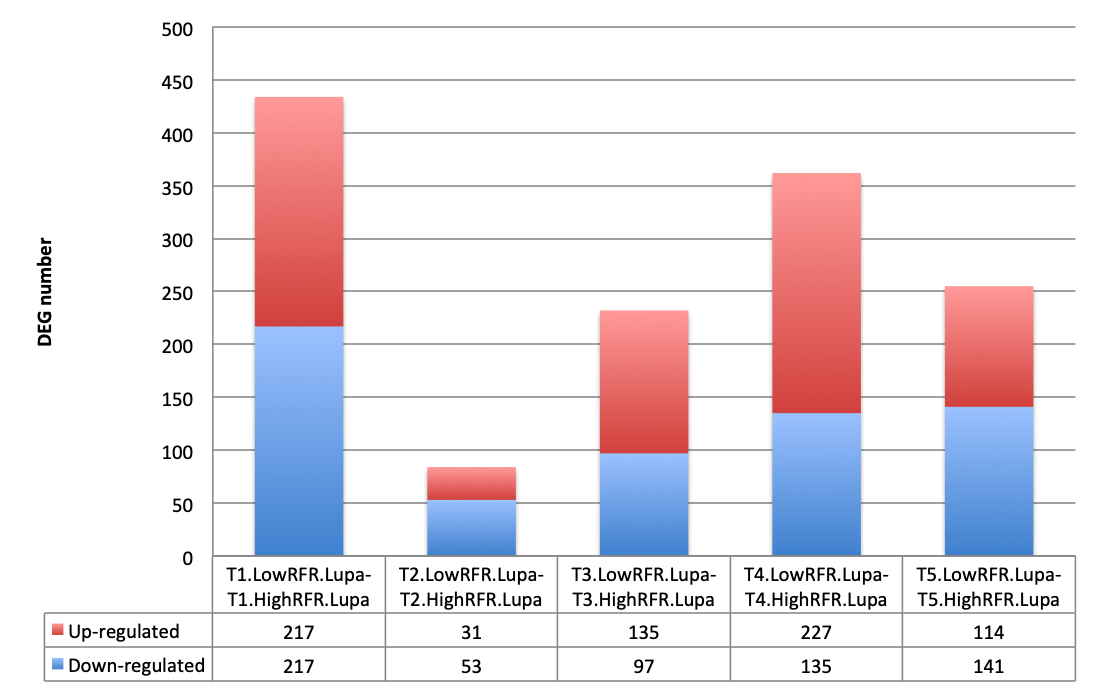

**A**

**B**

Fig. S3. Bar plots shown numbers of up-regulated and down-regulated genes from the contrast sets between low R/FR and high R/FR of five different growth stages for both *L. orientalis* BGE016880 (A) and *L. culinaris* cv. Lupa (B).

Supplementary Tables

| Gene | Primer-F | Primer-R |
| --- | --- | --- |
| *LcAGL6/13d* | CAATACAGTTCCACTGACTTGAAC | CTCCTGGTATAAACCCTGTGAC |
| *LcELF4a* | GAAGATCTTTGCTGTTGATGACAG | TCTAGAATGCCTGATAACATGGTG |
| *LcLWD1* | TAGGATCAAGCCCACCTTCCA | CTAGTGTGAATGCGATTGCGTG |
| *LcFTa1* | CATTGGTTGGTGACCGATATTCC | CCTTTGTCTACACTGCTGACGA |
| *LcFTb1* | CTCTGATTATGGTGAATCCTGATG | CATAGCTCACTATCTCTTGCCCT |
| *LcFTb2* | GTTACCATCCAATGAAGGTATTCC | GGTGAGCTCAAACCATCTCAAG |
| *LcFTc* | TGGTCGATCCTCATGTTGTAGG | TTGGACGATTAACCAATTGTGAGG |
| Actin | CTTTGCAATCCACATCTGTTGGAA | GCTTTGGCACCAAGTAGCATGA |

Table S1. Primer list used for verification of flower DEGs through RT-qPCR. Primers for qPCR were designed based on the consensus transcripts between *L. culinaris* cv. Lupa and *L. orientalis* BGE 016880 when aligned with annotated gene regions within the *L. culinaris* cv. CDC Redberry genome assembly (version 2.0).

| Linkage groups (LGs) | Markers mapped | Map length covered (cM) | Average inter-marker distance (cM) | Maximum Marker Distance (cM) |
| --- | --- | --- | --- | --- |
| LG1 | 496 | 705.6 | 1.4 | 8.2 |
| LG2 | 958 | 1444.2 | 1.5 | 8.3 |
| LG3 | 702 | 986.1 | 1.4 | 8.0 |
| LG4 | 730 | 1058.6 | 1.5 | 8.3 |
| LG5 | 590 | 864.7 | 1.5 | 9.4 |
| LG6 | 597 | 864.1 | 1.4 | 8.6 |
| Total | 4073 | 5923.3 | 1.5 | 9.4 |

Table S2. Detailed characteristics of the genetic linkage map of a lentil interspecific recombinant inbred line population developed from *L. culinaris* cv. Lupa and *L. orientalis* BGE 016880.

|  | *L. orientalis* BGE 016880 | *L. culinaris* cv. Lupa |
| --- | --- | --- |
| Total transcripts | 138108 | 135723 |
| Total genes | 78707 | 76595 |
| Percent GC (%) | 37.98 | 37.99 |
| Contig N50 | 2255 | 2252 |
| Median contig length | 845 | 877 |

Table S3. Statistics of *de novo* assembled Transcriptomes for both *L. orientalis* BGE 016880 and *L. culinaris* cv. Lupa. Trinity was used for *de novo* assembly of the transcriptomes and the statistics was generated using TrinityStats.

Supplementary Methods

Differential gene expression analysis of RNAseq data

3D RNA-seq pipeline (Guo et al., 2019; Calixto et al., 2018) was used for the analysis of differential gene and transcript expression within species. Details as follow: The RNAseq data had 10 factor groups (T1.HighRFR, T2.HighRFR, T3.HighRFR, T4.HighRFR, T5.HighRFR, T1.LowRFR, T2.LowRFR, T3.LowRFR, T4.LowRFR, T5.LowRFR) and each had 3 biological replicates for both *L. culinaris* cv. Lupa and *L. orientalis* BGE 016880 (30 samples each and 60 samples in total). Read counts and transcript per million reads (TPMs) were generated using tximport R package and lengthScaledTPM method (Soneson et al., 2016) with inputs of transcript quantifications from Salmon (Patro et al., 2017). Low expressed transcripts and genes were filtered based on analyzing the data mean-variance trend. Expressed transcripts were determined as which had ≥ 1 of the 30 samples with count per million reads (CPM) ≥ 1. A gene was expressed if any of its transcripts with the above criteria was expressed. The TMM method was used to normalize the gene and transcript read counts to log_2_-CPM (Bullard et al., 2010). RUVSeq R package with approach RUVr was used to estimate the batch effects (Risso et al., 2014). Limma R package was used for differential expression comparison (Ritchie et al., 2015; Law et al., 2014). To compare the expression changes between low R/FR light quality environment and high R/FR light quality environment from five different growth stages, the contrast groups were set as T1.LowRFR-T1.HighRFR, T2.LowRFR-T2.HighRFR, T3.LowRFR-T3.HighRFR, T4.LowRFR-T4.HighRFR, and T5.LowRFR-T5.HighRFR for both *L. culinaris* cv. Lupa and *L. orientalis* BGE 016880. To compare the expression changes between low R/FR light quality environment and high R/FR light quality environment, the contrast group was set as (T1+T2+T3+T4+T5). LowRFR/5 - (T1+T2+T3+T4+T5). HighRFR/5 for both *L. culinaris* cv. Lupa and *L. orientalis* BGE 016880. For DE genes, the 𝑙𝑜𝑔_2_ fold change (Log_2_FC) of gene abundance was calculated based on contrast groups and significance of expression changes were determined using t-test. P-values of multiple testing were adjusted with BENJAMINI HOCHBERG (BH) procedure to correct false discovery rate (FDR) (Benjamini and Yekutieli, 2001). A gene/transcript was significantly differentially expressed in a contrast group if it had adjusted p-value < 0.05 and |Log_2_FC| ≥ 1.

Quantitative Reverse Transcriptase –PCR (qRT-PCR) analysis

To confirm the reliability of RNAseq results, representative flowering DEGs in the QTL regions, flower genes that were not DEGs but within the QTL regions, as well as flowering DEGs that were not in any QTL regions, were selected for qRT-PCR validation. Total RNA used for qRT-PCR analysis was isolated, quantified and qualified using the same procedures as those that were used for RNAseq study. SuperScript^TM^ IV First-Strand Synthesis System (Thermo Fisher Scientific Inc, Waltham, MA, USA) was used to synthesize cDNA from total RNA. Primers for qPCR were designed based on the consensus transcripts between *L. culinaris* cv. Lupa and *L. orientalis* BGE 016880 when aligned with the annotated gene regions within the *L. culinaris* cv. CDC Redberry genome assembly (version 2.0) visualized using Integrative Genomics Viewer (Robinson et al., 2011). The sequences of the primers used in qPCR were listed in supplemental information (Supplementary Table 6). qPCR reactions were set up using FAST SYBR® Green qPCR Master Mix (Applied Biosystems, Foster City, CA, USA) and amplifications were performed on a BioRad CFX384^TM^ Real-Time PCR system (Bio-Rad Laboratories, Inc., Hercules, CA, USA). qPCR condition used was as following: 95°C initial denaturation for 15 sec, followed by 40 cycles of 95°C denaturation for 3 sec plus primer annealing/extension at 60°C for 30 sec. Post-amplification melting-curve analyses were included to check for primer-dimer artifacts and reaction specificity (one cycle of 95°C for 15 sec, 65°C for 5 sec with a gradual increase in temperature (0.5°C/ 5 sec) to 95°C). Actin was selected as the reference gene due to its relative consistent stable expressions across different stages as well as different light quality environments from previous transcriptomic analysis. To derive the relative expression value, the delta-delta CT method was adopted and samples from T1 stage at low R/FR light quality environment were used as reference samples (Kenneth et al., 2001).
