## Supplementry File 6 for "Genetic and gene expression analysis of flowering time regulation by light quality in lentil"

```
#!/usr/bin/perl

use strict;
use warnings;
use Data::Dumper;

my $SAM_file = $ARGV[0];
my $deg_list = $ARGV[1];
my $usage = "./convert_TRINITY_DEG_list.pl input.SAM input.DEG_list";

unless (defined $SAM_file) {
        die "ERROR: Either a SAM file wasn't specified or it can't be
found. Usage: $usage";
}
unless (defined $deg_list) {
        die "ERROR: Either a DEG list wasn't specified or it could not
be found. Usage: $usage";
}

my @splitline;
my @splitread;
my %readnames;
my ( $readname, $gene, $trinityname );

# Store our read names from the SAM file into a hash
# Assumes a non-header SAM file
open SAMFILE, $SAM_file or die "ERROR: Unable to open $SAM_file\n";

while (my $alignment = <SAMFILE>) {

        chomp $alignment;
        @splitline = split(/\t/, $alignment);
        $readname = $splitline[0];
        # Alter the readname since the DEG list leaves off the "_i#"
        $readname =~ s/_i[0-9]*$//g;
        $trinityname = $readname;
        @splitread = split(/_/, $readname);
        $gene = $splitread[0];
        $trinityname =~ s/$gene/TRINITY/;
        ## There might already be a sequence ID with this name if the
read aligned to more than one gene,
        ## so treat this as a hash of arrays
        ## Since the DEG format is less specific for its Trinity
names, make sure we do exclude if we have this exact gene/TRINITY name
combo already
        unless (exists $readnames{$trinityname} && (grep /$gene/,
@{$readnames{$trinityname}})) {
                push @{ $readnames{$trinityname} }, $gene;
        }
}
```

```
close SAMFILE;

my $genename;

# Loop through the DEG list IDs and change any target names that were
in the SAM file
open DEG, $deg_list or die "ERROR: Unable to open $deg_list\n";

while (my $line = <DEG>) {
	foreach my $key (keys %readnames) {
		if ( $line =~ /$key\s/ ) {
			my $newtarget;
			foreach $genename ( @{$readnames{$key}} ) {
				$newtarget .= $genename . "_";
			}
			$line =~ s/TRINITY_/$newtarget/g;
			last;
		}
	}
	print $line;
}

close DEG;
```
